# Whole genomes reveal how Andean climate history shapes genetic diversity and modern conservation risk in South American pumas

**DOI:** 10.64898/2026.02.19.706801

**Authors:** Daniel E. Chavez, Chiara Correa-Zanotti, Carolina Sáenz, Lauren Ong, Nicole Ormaza, David Mora, María B. Cabezas, Alexander Medina, Robert K. Wayne, Tuyen Ong, Rebecca Zug

**Affiliations:** Centro de Investigación de la Biodiversidad y Cambio Climático (BioCamb), Facultad de Ciencias de Medio Ambiente, Universidad Tecnológica Indoamérica, Machala y Sabanilla, Quito, EC170301, Ecuador; Paris Lodron Universität Salzburg, Austria; Pampas Cat Working Group, Piura, 20000, Perú; Hospital de Fauna Silvestre Tueri, Instituto iBIOTROP, Universidad San Francisco de Quito (USFQ), Quito, Ecuador; Concord Academy, 166 Main St, Concord, MA 01742, United States; Fundación Zoológica del Ecuador. Pasaje Pircabamba y Rumichupa, Guayllabamba 170209, Ecuador; Departamento de Investigación y Biología, Mashpi Lodge, Pacto, Ecuador; Department of Ecology and Evolutionary Biology, University of California, Los Angeles, Los Angeles, California 90095, USA; Ring Therapeutics, Cambridge, MA 02139, USA; Laboratorio de Carnívoros, Colegio de Ciencias Biológicas y Ambientales, Universidad San Francisco de Quito (USFQ), Quito 170901

## Abstract

Climatic oscillations in the Andes have repeatedly reshaped habitats over millions of years, yet their long-term genomic consequences for wide-ranging carnivores remain unclear. We generated whole-genome sequences from pumas (*Puma concolor*) across ecologically distinct regions of Ecuador to test how paleoclimate shaped population structure, demography, and genetic load. We show that northwestern forest pumas persisted in long-term isolation within humid refugia, whereas northern Andean and southern Pacific populations reconnected intermittently during warm interglacial periods. Southern coastal pumas maintained persistently small effective population sizes, leading to elevated runs of homozygosity and increased burdens of homozygous loss-of-function variants. In contrast, northern populations historically remained larger but exhibit early signs of inbreeding in one individual, marked by long runs of homozygosity and a kinked tail phenotype. Our findings indicate that recent fragmentation may be disrupting historical connectivity. Restoring corridors around the western foothills could reestablish gene flow and reduce inbreeding risk, while targeted genetic rescue may support chronically isolated southern populations. By integrating paleoclimate history with genome-wide data, we provide a framework for region-specific conservation strategies that balance connectivity restoration with the preservation of local adaptation.

## Introduction

The puma (*Puma concolor*) is one of the most widely distributed terrestrial mammals in the Western Hemisphere, ranging from Canada to Patagonia and occupying a broad diversity of habitats, from temperate montane systems to tropical lowlands and Andean forests (Sunquist & Sunquist, 2002). This extensive ecological breadth suggests a strong capacity for dispersal and local adaptation, yet it also implies that populations may respond heterogeneously to historical and contemporary environmental pressures (Nowell & Jackson, 1996).

South America represents the evolutionary and demographic core of the puma lineage (Saremi et al., 2019). Fossil and genomic evidence indicate a South American origin followed by a colonization of North America around 200 kya, resulting in a deeper evolutionary history in southern populations (Chimento & Dondas, 2018; Culver et al., 2000; Matte et al., 2013; Saremi et al., 2019). This history implies that pumas in South America experienced the full spectrum of major climatic dynamics that have shaped most Neotropical biodiversity during the last three million years (Berta, 1987; Chavez et al., 2022; Escobar et al., 2021; Gentry, 1982; León et al., 2024; van der Hammen, 1974). In the Andean region, Pleistocene glacial–interglacial cycles drove repeated contractions and expansions of forested habitats, creating periods of population isolation and secondary contact across geographic gradients (Cossios et al., 2009; Escobar et al., 2021; León et al., 2024; Matte et al., 2013; Webb, 1978). Consequently, pumas offer an informative model for investigating how climatic cycles interacted with dispersal ability to shape patterns of contemporary genomic diversity in the Andes Mountains.

Although pumas are currently classified as Least Concern (LC) under the IUCN Red List for Threatened species across much of South America (Nielsen et al., 2015), increasing anthropogenic pressures, including habitat fragmentation (Figure 1), hunting, and extractive activities, are likely affecting populations unevenly across the continent (Armijos-Arostegui et al., 2026; Castilho et al., 2011; Saranholi et al., 2017). Most recent genetic analyses of South American puma populations relied on mitochondrial DNA and microsatellites, which, despite broad geographic sampling, provide limited resolution for reconstructing demographic history and offer little insight into the functional consequences of genetic variation (Caragiulo et al., 2014; Castilho et al., 2011; Culver et al., 2000; Elbroch et al., 2024; Mac Allister et al., 2024; Matte et al., 2013; Saranholi et al., 2017). Whole-genome sequencing overcomes these limitations by capturing genome-wide variation, enabling more precise inference of population history while directly linking genetic diversity to functional regions affecting fitness (Aguilar-Gómez et al., 2025; Gustafson et al., 2022; Ochoa et al., 2022; Saremi et al., 2019). In particular, analyses of runs of homozygosity (ROH) provide a powerful framework for detecting long-term isolation and recent inbreeding, quantifying deleterious genetic load, and identifying populations at risk before demographic declines become evident (Kyriazis et al., 2025; Robinson et al., 2023).

**Figure 1.**
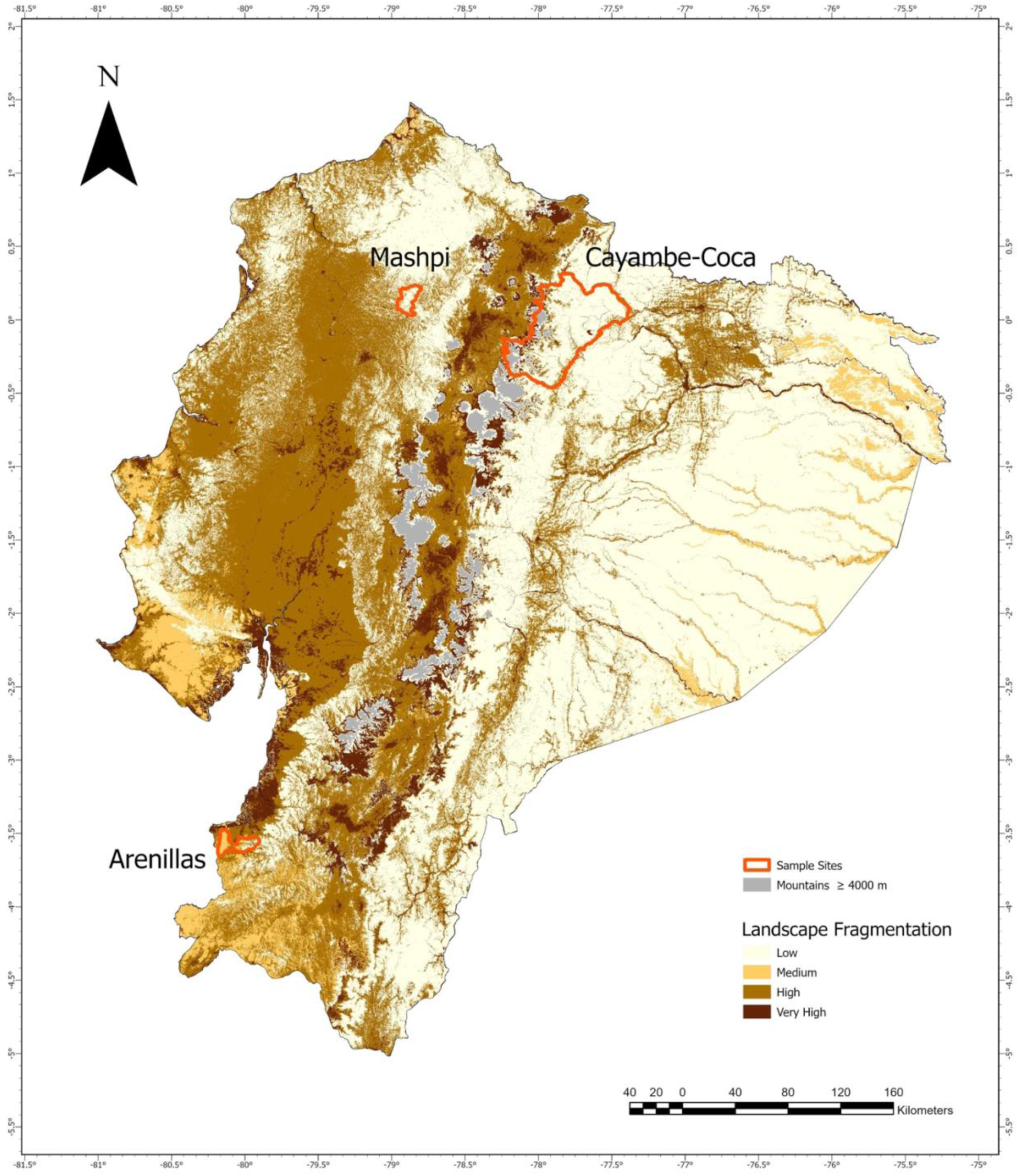
Landscape fragmentation and sampling locations of puma populations across Ecuador. Shaded areas represent increasing levels of landscape fragmentation from low to very high, with regions above 4,000 m elevation shown in gray. Red outlines indicate sampling areas for the Mashpi (northwestern Andes), Cayambe–Coca (northern Andes), and Arenillas (southern Pacific coast) populations.

Here, we present a whole-genome analysis of pumas from multiple geographic sites across the Ecuadorian Andes and the Pacific coast to investigate how evolutionary history across ecologically distinct landscapes has shaped contemporary genetic diversity. To accomplish this goal, we expanded genomic sampling beyond the two genomes previously studied from the Brazilian Amazon (Saremi et al., 2019) and assessed contemporary conservation status by integrating genome-wide variation, runs of homozygosity, and demographic inference.

## Results

### Population Structure and Admixture

Our principal component analysis of 20,760 SNPs revealed clear population structure, with individuals clustering by locality. Kinship analysis identified one sample from Mashpi, a cloud forest reserve in the western foothills of the Andes (Figure 1), with a kinship coefficient >0.2 (Figure S1), consistent with a sibling pair, which was subsequently excluded to minimize related-individual bias. We observed that pumas from the Arenillas population, an area located on the Pacific coast of southern Ecuador (Figure 1), separated clearly from the two Ecuadorian populations along PC2, which explained 26% of the genetic variance and distinguished southern from northern Ecuador (Figure 2b). One population was from Cayambe-Coca National Park, a heterogeneous landscape spanning roughly 4,000 km^2^ located at northern Ecuador (Figure 1). The second population was Mashpi. The PC1, also explaining 26% of the variance, further differentiated the Arenillas population and the two individuals from Cayambe-Coca from the remaining Cayambe-Coca and Mashpi individuals (Figure 2b). Overall, Arenillas emerged as the most genetically divergent of our sampled Ecuadorian populations. In contrast, Cayambe-Coca and Mashpi clustered closely along both axes, indicating relatively weak differentiation between these northern populations.

**Figure 2.**
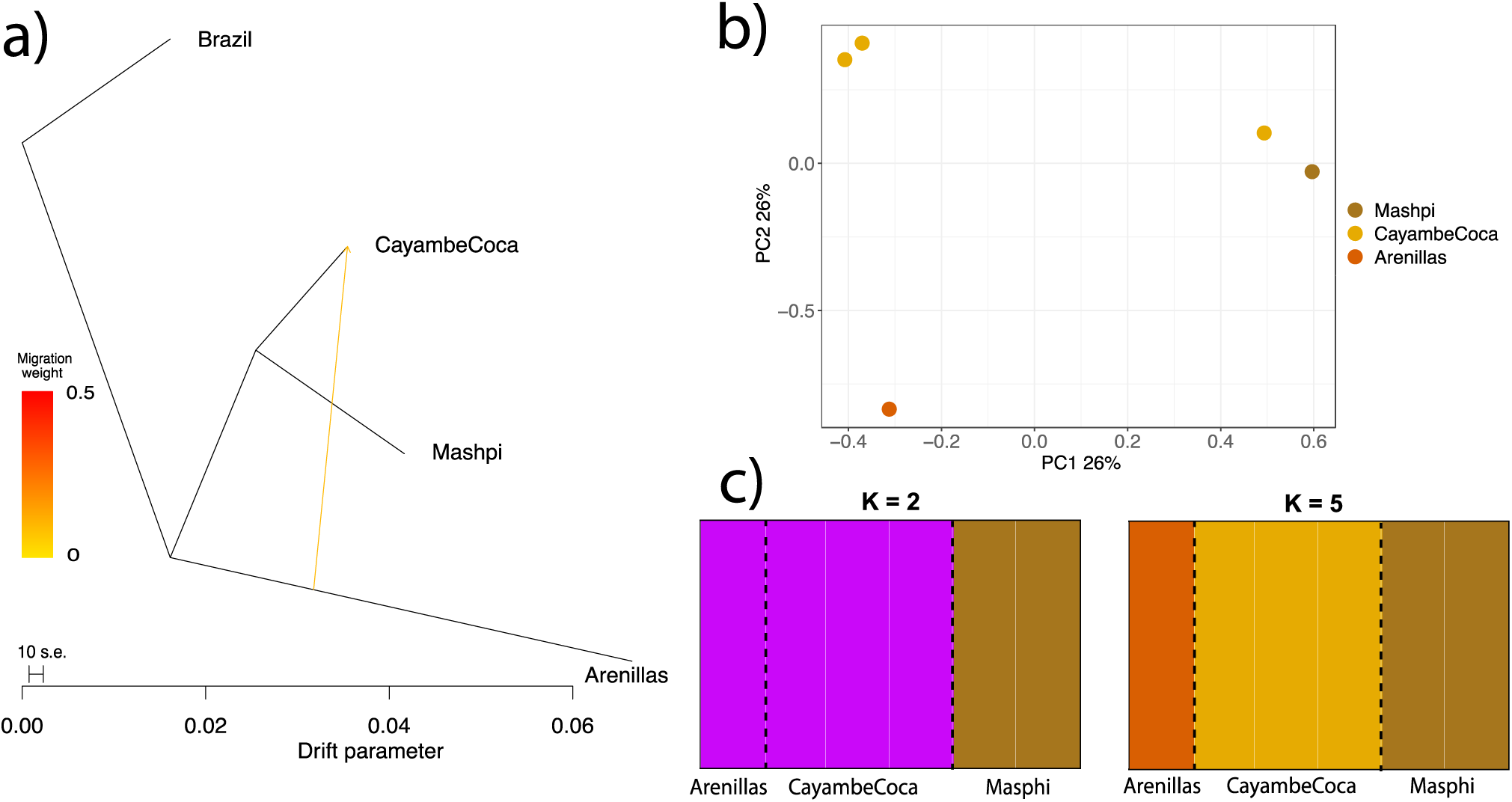
Evolutionary history of Andean pumas. **a)** TreeMix analysis showing genetic drift (x-axis) among Ecuadorian puma (*Puma concolor*) populations, using Brazil as the root population. The tree reveals clear differentiation among the Ecuadorian populations-Mashpi, Cayambe-Coca, and Arenillas-with the latter population exhibiting the highest drift values or differentiation. One migration event is inferred (yellow edge), suggesting limited gene flow between the Arenillas and Cayambe-Coca populations. **b)** Principal component analysis (PCA) based on 20, 760 independent SNPs. PC1 explains the 26% of the variation and separates the Arenillas population and two individuals from Cayambe-Coca from other Ecuadorian populations (Mashpi), as well as one individual from Cayambe-Coca. PC2, which explains the 26% of the variation, differentiates the Arenillas population from the rest of Ecuadorian populations. **c)** fastSTRUCTURE analysis for K =2 and K = 5. Each vertical bar represents an individual, and colors denote inferred genetic ancestry. At K=2 The results highlight greater ancestry similarity between the Arenillas and Cayambe-Coca populations.

Genetic-connectivity analyses grouped Mashpi and Cayambe-Coca into a closely related clade, directly supporting the population structure revealed by PCA (Figure 2a). Consistent with the PCA results, the Mashpi and Cayambe-Coca populations formed a closely related clade (Figure 2a). In contrast, the Arenillas population exhibited a long branch with a drift value of 0.07, approximately four times greater than that observed in the other Ecuadorian populations (Figure 2a). Notably, TreeMix detected evidence of admixture between the Arenillas and Cayambe-Coca populations, suggesting that pumas historically occupied a broader and more connected distribution along the Andes (Figure 2a).

Genetic clustering analyses supported a consistent population structure across models, with K values from 1 to 10 yielding similar marginal likelihoods. Consistent with the signal of gene flow inferred from the TreeMix results, at K=2 individuals from the Arenillas and Cayambe-Coca populations grouped into a single genetic cluster. In contrast, the Mashpi population formed a distinct cluster (Figure 2c). This clustering pattern remained stable from K = 2 through K = 4. At K = 5, individuals from Cayambe-Coca, Mashpi, and Arenillas formed three independent clusters (Figure 2c). The subdivision observed at K = 5 may indicate more recent population separation, potentially driven by habitat fragmentation (Figure 1) and reduced connectivity across the landscape. Together, the similarity between the Arenillas and Cayambe-Coca at K = 2 (Figure 2) suggests historical gene flow between these populations, whereas the additional subdivision observed at K = 5 may reflect more recent population differentiation driven by recent separation or habitat fragmentation (Figure 1 and Figure 2).

### Genetic Diversity

Patterns of genome-wide heterozygosity revealed substantial regional variation, consistent with the influence of past demographic processes (Figure 3). Across all Ecuadorian populations, individuals exhibited regions of high heterozygosity along most autosomes (Figure 3a). However, sliding-window heterozygosity was consistently lower in the Arenillas population, suggesting a long-term smaller effective population size.

**Figure 3.**
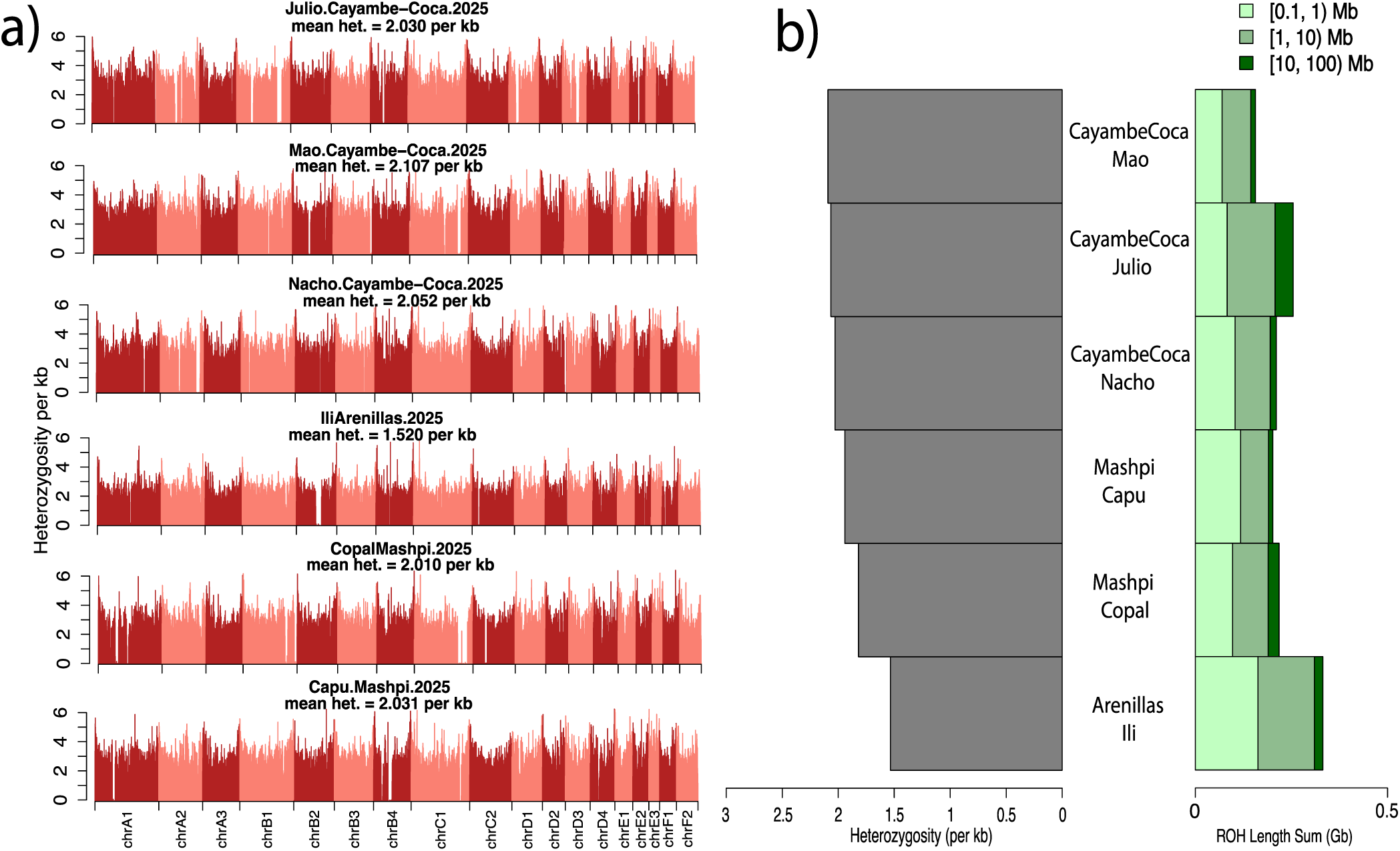
Genome-wide diversity and distribution of runs of homozygosity (ROH) in Ecuadorian pumas. **a)** Genome-wide heterozygosity profiles for individuals from Cayambe-Coca, Mashpi, and Arenillas. Panels show per-site heterozygosity across the autosomal genome, with mean heterozygosity values reported for each individual. Chromosomes are shown sequentially (chrA1–chrF2), illustrating stretches of variable diversity across the genome. **b)** Summed lengths of ROH across three size classes per individual: short ROH (0.1–1 Mb), medium ROH (1–10 Mb), and long ROH (10–100 Mb).

In addition, we identified several multi-megabase regions depleted of heterozygosity. These genomic regions were most pronounced in one individual from Arenillas (e.g., chromosome B2 in Figure 3a), but were also observed in a single individual from Cayambe-Coca (Julio; e.g., chromosomes B1, C2, and D4) and in the two individuals from Mashpi (Copal on chromosome C1 and Capu on chromosome B4). These extended homozygous regions may be consistent with recent inbreeding events. Despite these localized patterns, genome-wide heterozygosity was broadly similar across populations, ranging from 1.5 heterozygous sites per kb in Arenillas to 2.1 in Cayambe-Coca (Figure 3a, Table S1).

Runs of homozygosity highlighted genomic regions with depleted genetic variation. The highest burden of long ROH was observed in Julio, an individual from Cayambe-Coca (Figure 3b). Julio exhibited approximately 47 Mb of long ROH, representing ~2% of the genome, which is more than twice the Ecuadorian population average of 20 Mb (Table S1). Importantly, this individual also displayed a kinked tail (see <u>Supplementary Video</u>) a phenotype frequently associated with recent inbreeding (Gustafson et al., 2022; Huffmeyer et al., 2022; Johnson et al., 2010). Additionally, Julio exhibited elevated levels of medium-sized ROH (~124 Mb; ~5% of the genome), exceeding those observed in most other individuals, except Ili from the Arenillas population, who showed the highest amount of medium-sized ROH (146 Mb; ~6% of the genome). This individual also showed extensive short ROH, totaling 162 Mb (~7% of the genome; Figures 3b and Table S1). The predominance of short and medium ROH in Ili is consistent with long-term persistence at a small effective population size rather than with exclusively recent inbreeding.

Finally, individuals from the Mashpi and Cayambe-Coca populations, with higher genome-wide heterozygosity (~0.8 heterozygous sites per kb), exhibited few long ROH (>2 Mb), ranging from 1 to 2 Mb and representing less than 1% of the genome (Figure 3b and Table S1). Overall, these results indicate that recent population declines and inbreeding have been most pronounced in the Cayambe-Coca region of northern Ecuador (see Julio in Figure 3b), whereas the southern Ecuadorian population, Arenillas, shows genomic signatures consistent with a long-term small population size (Figure 3b).

### Ancestral Climatic Reconstruction

Consistent with the genomic evidence of episodic connectivity and long-term isolation, our paleoclimatic reconstruction revealed temporal convergence between Cayambe-Coca and Arenillas under warm conditions and persistent climatic separation of Mashpi (Figure 4). In particular, during the Mid-Pliocene Warm Period (mPWP; ~3.2 Ma), and to a lesser degree during the Last Interglacial (LIG ~ 130kya), climatic differences between Cayambe-Coca and Arenillas was markedly reduced in comparison with a much colder phase near the end of the last ice age between 17 kya and 14.7 kya during the Heinrich Stadial 1 (HS1; Figures S2–S4). This means that environmental conditions in northern and southern regions became much more similar than during Pleistocene cold periods, potentially reducing ecological barriers between populations. This climatic convergence during warm periods is consistent with signals of historical gene flow between these populations inferred from TreeMix and STRUCTURE analyses (Figure 2), which indicate past connectivity despite their present-day geographic separation.

**Figure 4.**
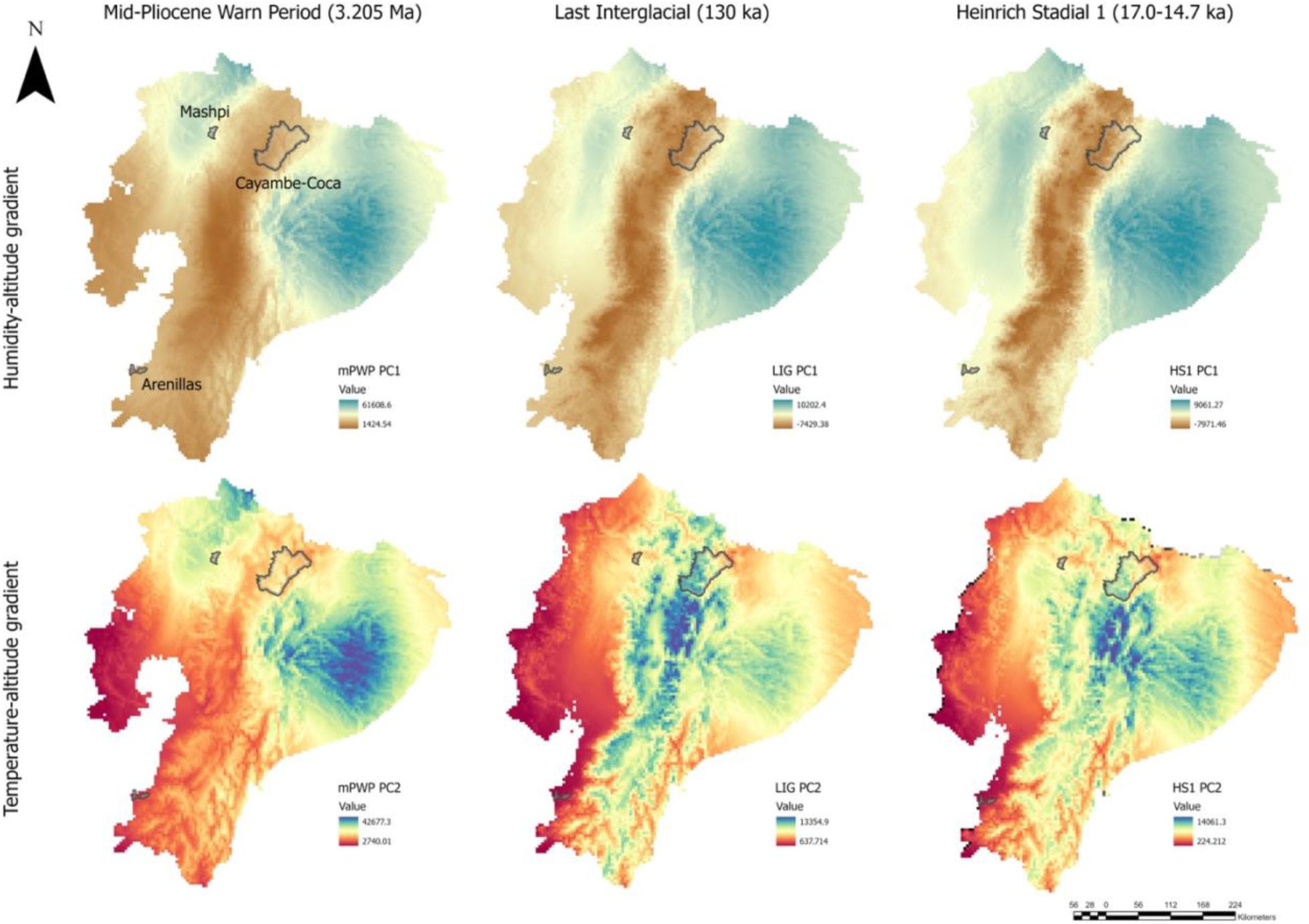
Spatial projections of principal climatic gradients across paleoclimatic periods. The top row represents humidity–altitude gradients (see PC1 in Figures S2–S4). The bottom row represents temperature–altitude gradients (see PC2 in Figures S2–S4). Polygons for sampled populations for Mashpi, Cayambe-Coca, and Arenillas are overlaid. Panels illustrate climatic variation across the Mid-Pliocene Warm Period (3.205 Ma), Last Interglacial (~130 ka), and Heinrich Stadial 1 (~17–14.7 ka).

In contrast, during the major cold phase near the end of the last ice age, between 17kya and 14.7 kya (HS1), Cayambe-Coca and Arenillas experienced very different climatic conditions (Figure 4). Specifically, Cayambe-Coca was associated with cold, high-elevation conditions, whereas Arenillas occupied warmer, drier environments (Figures S2–S4). Notably, climatic distances between these populations were large in both intervals, consistent with the population divergence inferred from TreeMix.

Lastly, Mashpi consistently experienced different environmental conditions across all periods examined (Figure 4 and Figures S2–S4). This persistent difference matches genomic evidence showing long-term isolation from other puma populations.

### Demographic History

We observed contrasting demographic trajectories over the past 500,000 years for pumas (Figure 5). We first examined the southern population from Arenillas, which previously showed low genetic diversity and long runs of homozygosity (ROH). This population exhibited a relatively stable but consistently low Ne of approximately 80,000 for most of its history, followed by a pronounced decline to ~10,000 during the past 20,000 years. In contrast, the northern populations (Cayambe-Coca and Mashpi) maintained larger and stable effective population sizes of approximately 100,000 until the late Pleistocene. Both populations experienced a sharp reduction in Ne over the past 20,000 years, reaching an estimated present-day population size of ~20,000 (Figure 5).

**Figure 5.**
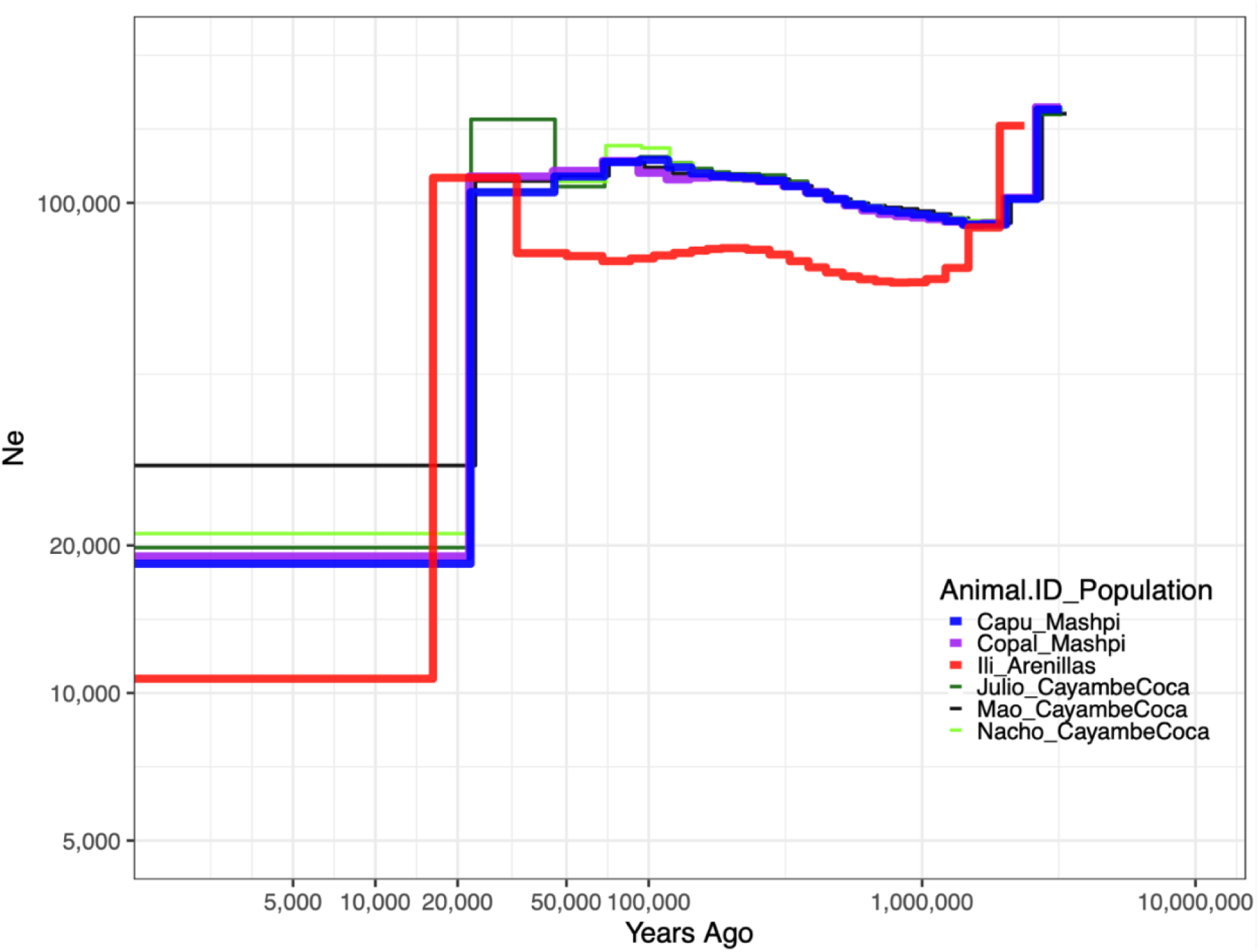
Demographic history of the Ecuadorian puma inferred using MSMC. MSMC-inferred demographic trajectories for six pumas (*Puma concolor*) sampled from Cayambe-Coca (Julio, Mao, Nacho), Mashpi (Capu, Copal), and Arenillas (Ili). The y-axis represents effective population size (Ne) inferred from inverse instantaneous coalescent rates (IICR) scaled by 2µ, and the x-axis shows time in years before present.

Overall, MSMC-derived demographic trajectories refine the coarse-scale patterns inferred from genomic diversity metrics (Figure 5). Together, these results suggest that the low genetic diversity and extended ROH observed in the Arenillas population reflect a long-term small population, whereas the northern Andean populations historically persisted at substantially larger sizes.

### Deleterious Variation

We investigate the effects of demographic history on deleterious variation by quantifying the accumulation of putatively deleterious mutations. We observed that Pumas from the Arenillas population exhibited the highest proportion of homozygous derived loss-of-function (LOF) genotypes, with approximately 25% of alleles falling into this deleterious category. This represents an increase of about three percent more than other Ecuadorian populations, which averaged ~22% of alleles classified as LOF (Figure 6; Table S2). In contrast to this pronounced enrichment of LOF variants, Ili, from Arenillas, showed only a modest increase (~1%) in benign homozygous derived genotypes, with missense mutations accounting for 18% of alleles compared to an average of 17% in other populations (Table S2). Together, these patterns indicate that the elevated burden of damaging variants in the Arenillas population is driven primarily by long-term reductions in effective population size. This interpretation is consistent with theoretical expectations that selection is less efficient in small populations, allowing mildly deleterious mutations to persist or reach fixation (Charlesworth & Willis, 2009).

**Figure 6.**
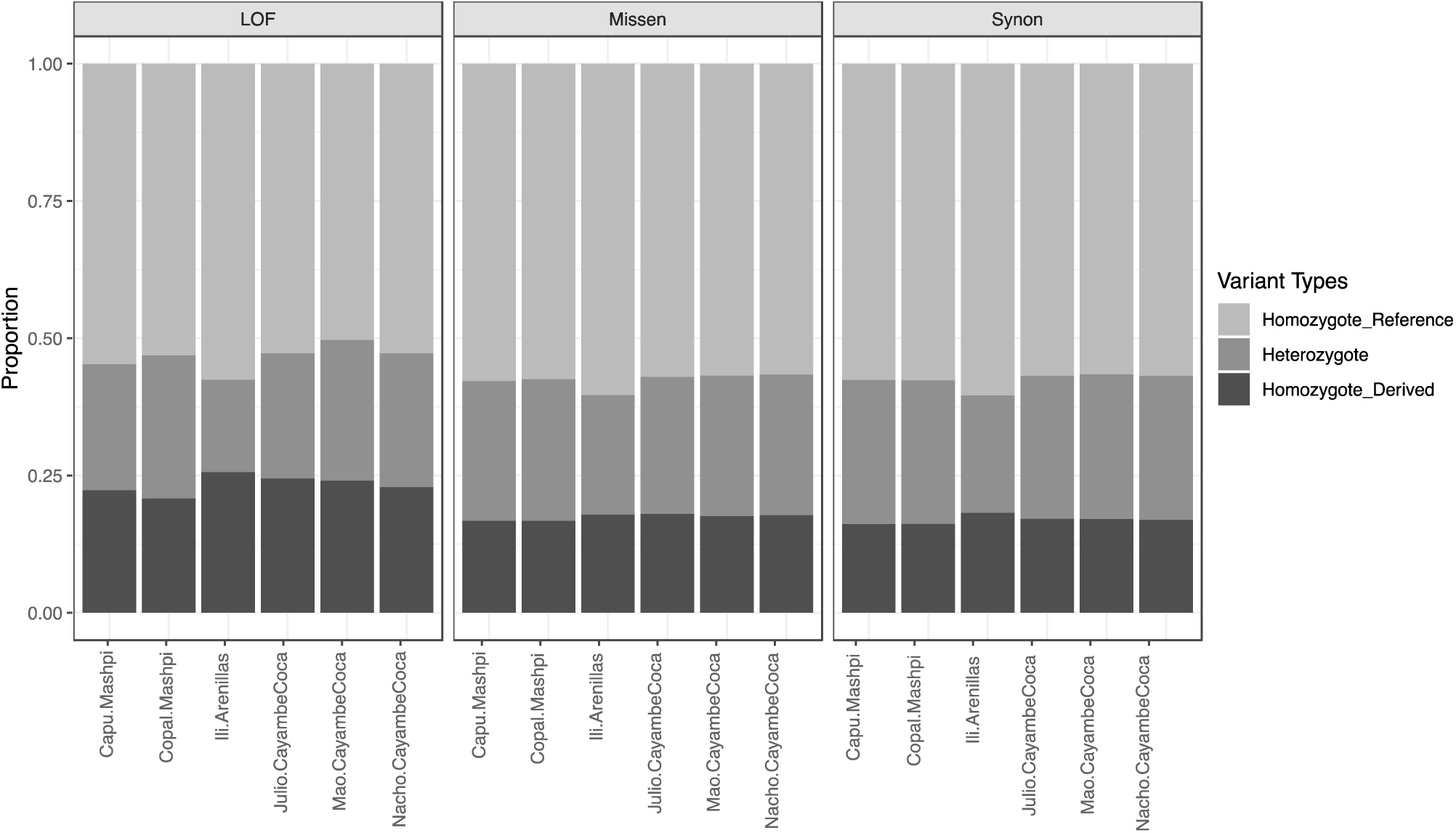
Proportion of derived alleles across functional categories in Ecuadorian pumas. Proportions of ancestral and derived genotypes across three classes of genetic variants: loss-of-function (LOF), missense mutations (Missen), and synonymous mutations (Synon) in six Ecuadorian pumas from Mashpi (Capu, Copal), Cayambe-Coca (Julio, Mao, Nacho), and Arenillas (Ili). Bars represent the proportion of homozygous-Refrence, heterozygous, and homozygous-derived genotypes for each individual. Across the most damaging variant category, LOF, the Arenillas genome displays a higher proportion of homozygous-derived alleles relative to Andean individuals, consistent with reduced heterozygosity and a longer-term smaller effective population size in this southern population.

In contrast to the long-term signal observed in the Arenillas population, we detected evidence of more recent demographic effects in northern Ecuador. Julio, from Cayambe-Coca, exhibited the highest proportion of missense mutations among all sampled pumas, with 18% of variants classified in this deleterious category (Figure 6, Table S2). By contrast, the proportion of synonymous (presumably benign) mutations in Julio was comparable to that observed in Mashpi and Cayambe-Coca populations, indicating that the excess variation is concentrated in damaging categories. Combined with the high burden of long ROH and the presence of a kinked tail phenotype, this pattern supports the interpretation that the accumulation of damaging missense variants in Julio may reflect recent population decline and inbreeding.

## Discussion

### Historic connectivity and isolation of puma populations across the Andes

We evaluate admixture to better understand ancestral distribution and connectivity of the puma population in Ecuador over evolutionary timescales. Our genomic analyses reveal that puma populations in the Andes do not mix freely (Figure 2). Instead, they show a pattern of long-term isolation in some regions and intermittent contact in others (Figure 4). These patterns reflect population dynamics guided by climatic isolations that operated from approximately 3.5 million years ago through the late Pleistocene around 14 kya (Gentry, 1982; Heine, 2000; Rodbell et al., 2009; van der Hammen, 1974; Webb, 1978).

One striking result is the persistent isolation of pumas in the northwestern Andean slopes (Figure 1 and Figure 4), where the Mashpi puma population occurs in the foothills, yet it shows stability and climatic similarity to a biogeographic region called the Chocó, a large and elongated humid tropical forest that runs along the Pacific coast from eastern Panama to northwestern Ecuador (Gentry, 1982; Pérez-Escobar et al., 2019). Our genome-wide analyses of principal components and admixture graphs placed the population of Mashpi as a distinct, non-admixed lineage. This was not expected for a highly mobile carnivore, especially considering that the closest population we sequenced, Cayambe-Coca, is only 100 km east (Figure 1). Notably, this distance is three times less than the recorded dispersal distances of juvenile pumas (Stoner et al., 2008). Importantly, this inference reflects deep-time population separation, from 3.5mya until 17kya, predating extensive human occupation of the inter-Andean valley, and therefore cannot be attributed to recent anthropogenic barriers. Therefore, the lack of gene flow could be related to the climate history in the Andes. Despite repeated glacial cycles, the western foothills remained wet. Cloud forests and rainforests persisted while surrounding habitats dried (Behling et al., 1998; Prance, 1982). Our paleoclimatic reconstructions further show that Mashpi remained climatically distinct during the last 3.5 million years (Figure 4). Previous studies have shown that humid refuges offered sanctuary for forest-dependent populations, such as Mashpi (Escobar et al., 2021). Our results indicate long-term environmental and genomic isolation of puma populations in the Ecuadorian Northwest.

In contrast, pumas from Arenillas in the south and Cayambe–Coca in the north bear genomic signatures of admixture despite being 470 km apart (Figure 1), which is more than four times the distance with respect to the distance between the two northern populations (Mashpi and Cayambe-Coca). We interpret this evidence of historical connectivity and secondary contact as the fingerprint of changing landscapes. During warmer, wetter interglacial intervals, savanna-like habitats expanded along the Andean route (Figure 4). These prolonged, contiguous landscapes would have opened corridors for movement (Berta, 1987; Heine, 2000; van der Hammen, 1974). For example, analyses of fossil records indicate that canids and felids migrated between northern and southern regions of the Andes via an Andean corridor that extended across the central Andes and reach the Pacific coast (Berta, 1987; Webb, 1978). Our climatic reconstruction provides independent support for this interpretation. During two warm periods about 3.2 mya and 130 kya, northern Andean and southern coastal regions experienced similar climatic conditions (Figure 4). Such climatic convergence may correspond to periods when pumas expanded across broad geographic regions, while population structure was due reinforced by isolation during colder intervals. This pattern has been repeatedly documented in genetic studies showing widespread distribution alongside strong regional differentiation and subspecies-level divergence across the Americas (Caragiulo et al., 2014; Culver et al., 2000; Matte et al., 2013).

When the warm periods that maintained these corridors ended and the climate cooled, glacier expansion and changes in the habitat disrupted landscape connectivity (Heine, 2000; Rodbell et al., 2009). As a result, puma populations became isolated, explaining the genetic differentiation observed between the Arenillas population in the south and the Cayambe–Coca population in the north, despite evidence of historical admixture. This suggests that along the latitudinal axis, climate swings repeatedly opened and closed pathways for contact (Berta, 1987; Cossios et al., 2009; Escobar et al., 2021; Gentry, 1982; Webb, 1978). Our climatic reconstruction was consistent with these scenarios. Notably, the strong climatic compartmentalization during Pleistocene intervals, particularly during Heinrich Stadial 1 (17–14.7 ka), supports scenarios of environmental isolation that may have reinforced population divergence. These findings are consistent with broader phylogeographic models proposing that climatic oscillations act as major drivers of connectivity and divergence in montane systems (Ramírez-Barahona & Eguiarte, 2013; Turchetto-Zolet et al., 2013).

### Long-term demographic constraint in southern populations

Genomic patterns in the Arenillas puma population illustrate the consequences of long-term demographic constraint and persistently small population size rather than recent inbreeding. Individuals from the Arenillas population exhibit a relatively high proportion of medium-length ROH, a pattern indicative of ancient reductions in effective population size, as recombination over many generations progressively breaks down long ROH into shorter segments (Kyriazis et al., 2025). This genomic signal is consistent with the biogeographic context of southwestern Ecuador, which forms part of the Tumbesian seasonally dry forest region (Dodson & Gentry, 1991). This region is geographically restricted, approximately half the extent of the Chocó (Gentry, 1982), and highly fragmented across southwestern Ecuador and northwestern Peru (Dodson & Gentry, 1991). Despite its limited size, it represents a climatically distinct ecosystem that has persisted as a long-standing biogeographic unit through late Quaternary climate fluctuations (Heine, 2000; Webb, 1978).

Previous genomic studies of carnivores have reported reduced effective population size in the Tumbesian region (see Sechuran fox in Chavez et al., 2022), suggesting that long-term demographic constraint may be a common feature of carnivore populations inhabiting this biogeographic region (Berta, 1987; Webb, 2006; Webb, 1978). Similar to other puma populations in North America (Gustafson et al., 2022; Huffmeyer et al., 2022; Johnson et al., 2010; Saremi et al., 2019), such environmental differentiation and historical habitat restriction are expected to limit effective population size and amplify the effects of genetic drift, even in the presence of sporadic gene flow from northern Andean populations such as Cayambe–Coca. Consequently, Arenillas’ pumas harbor the highest proportion of homozygous derived loss-of-function variants with respect to other pumas analyzed in this study. This result is expected under a prolonged small effective population size, where genetic drift permits strongly deleterious alleles to persist. Collectively, these results indicate that genomic erosion in the Arenillas population reflects past demographic constraint associated with the Tumbesian region rather than the acute effects of recent inbreeding.

### Early Signals of Inbreeding or Locus-Specific Effects Underlying Tail Malformation

We evaluated demography, population structure, and genetic load to better understand the conservation status of the puma population in the Andes. Our genomic results show that one individual from Cayambe-Coca (Julio) exhibited the highest burden of long runs of homozygosity among all sampled pumas, along with a kinked tail phenotype commonly associated with recent inbreeding (Huffmeyer et al., 2022; Johnson et al., 2010; Ochoa et al., 2022; Roelke et al., 1993; Saremi et al., 2019). Despite this result, Julio does not show genome-wide patterns that are typically diagnostic of strong recent inbreeding (Aguilar-Gómez et al., 2025; Chavez et al., 2022; Robinson et al., 2019), such as a high proportion of long ROH that can reach 20% the genome or the characteristic indentation pattern in heterozygosity landscapes observed in inbred individuals (Chavez et al., 2019; Robinson et al., 2019). In addition, we did not observe a high burden of deleterious alleles in this individual. This apparent discrepancy suggests two alternative interpretations that warrant careful consideration.

One possibility is that Julio has experienced a degree of recent inbreeding (after only a few generations of inbreeding) that is sufficient to generate phenotypic consequences, even though long ROH account for only a modest fraction of the genome. A growing body of genomic studies has shown that extended ROH are among the strongest predictors of inbreeding depression because they increase the likelihood that recessive deleterious alleles are expressed in homozygous form (Aguilar-Gómez et al., 2025; Kyriazis et al., 2023; Kyriazis et al., 2021; Robinson et al., 2023). Importantly, morphological abnormalities such as distal tail kinks have been repeatedly documented in inbred puma populations across North America and are widely regarded as reliable phenotypic indicators of elevated genomic homozygosity (Gustafson et al., 2022; Huffmeyer et al., 2022; Johnson et al., 2010). Empirical evidence from puma (mountain lion) populations in Southern California and Florida demonstrates that kinked tails frequently co-occur with high ROH burdens and reduced reproductive fitness, including compromised sperm morphology and teratospermia (Huffmeyer et al., 2022). Under this interpretation, Julio’s kinked tail and elevated ROH burden may reflect localized or moderate inbreeding with potential fitness consequences that are not fully captured by genome-wide summaries of long ROH.

Alternatively, Julio may not be inbred at the genome-wide level, and the kinked tail phenotype could instead arise from homozygosity at one or a few specific loci involved in tail development. Previous genetic studies in mammals have demonstrated that skeletal and axial malformations can arise from homozygosity at a limited number of key developmental genes involved in caudal and vertebral patterning, even in otherwise outbred individuals (Buckley et al., 2020; Hytönen et al., 2009; Pollard et al., 2015; Xu et al., 2016). If one of Julio’s ROH overlaps genomic regions regulating vertebral patterning or tail morphogenesis, homozygosity at such loci could produce a kinked tail phenotype that phenocopies traits commonly associated with inbreeding. Under this scenario, the morphological abnormality would reflect locus-specific effects rather than genome-wide inbreeding, reconciling the presence of a kinked tail with the absence of extensive long ROH and typical heterozygosity patterns (Aguilar-Gómez et al., 2025; Robinson et al., 2019; Saremi et al., 2019).

### Genome-Based Conservation Strategies

Our results suggest that the Arenillas puma population in southern Ecuador has maintained a relatively small population size compared to other Ecuadorian populations from approximately 1 million years ago to the present. However, genome-wide heterozygosity patterns do not resemble those of natural populations that have experienced a prolonged history of extremely small effective population size (Chavez et al., 2022; Robinson et al., 2019). This suggests that efficient purging of deleterious alleles is unlikely to have occurred in the Arenillas population, a conclusion consistent with the elevated proportion of deleterious alleles observed in our study (Figure 6). Under this scenario, sporadic historical gene flow between pumas from Northern and southern populations was sufficient to maintain moderate levels of genetic diversity in the former population. Such gene flow may also have slowed the accumulation and expression of highly deleterious alleles, as shown in previous studies in wild populations (Dussex et al., 2025; Gustafson et al., 2022; Kyriazis et al., 2024). Several past episodes of admixture may have prevented the Arenillas population from reaching the high levels of homozygous deleterious variation typically observed in severely inbred populations.

Importantly, contemporary human-driven habitat fragmentation (Figure 1) has now eliminated the historic connection between the northern Cayambe-Coca population and the southern Arenillas populations (Berta, 1987; Heine, 2000; van der Hammen, 1974), effectively isolating the latter population and potentially accelerating drift-driven genomic erosion beyond historical levels. Forward-time genomic modeling studies show that once connectivity is lost, drift-driven accumulation of genetic load can accelerate rapidly in naturally occurring populations (Kyriazis et al., 2024), defining conditions under which alternative strategies such as genetic rescue become a plausible management option (Aguilar-Gómez et al., 2025; Johnson et al., 2010; Kyriazis et al., 2024). From a conservation perspective, the combination of long-term isolation and small population size suggests that the Arenillas population on the Pacific coast may be particularly vulnerable. As a result, this population could benefit from assisted gene flow using genetically compatible source populations to improve long-term persistence (Aguilar-Gómez et al., 2025; Fitzpatrick & Keller, 2015; Johnson et al., 2010; Ochoa et al., 2022).

In the case of the Mashpi population, these pumas represent an ecologically distinct lineage that has persisted in the northwestern forests with no detectable historical gene flow from other Andean puma populations, making the preservation of locally adapted variation a central conservation concern (Aguilar-Gómez et al., 2025). For Mashpi, long-term persistence under humid forest conditions suggests that assisted gene flow from climatically dissimilar populations such as Cayambe–Coca or Arenillas could dilute locally adapted haplotypes and reduce adaptive potential (Adams et al., 2011; Fitzpatrick & Keller, 2015; Johnson et al., 2010). For the Florida panther (puma), for example, genetic rescue produced strong demographic gains but also led to a reduction in the frequency of unique haplotypes (Aguilar-Gómez et al., 2025; Johnson et al., 2010; Ochoa et al., 2022). In this context, maintaining or restoring landscape connectivity among populations of the western foothills and the broader Chocó region, where environmental conditions and selective pressures are more similar, may be a more appropriate strategy to reduce inbreeding risk while minimizing the loss of locally adapted haplotypes.

Lastly, our results suggested that the pumas from Cayambe–Coca represented the largest population of pumas included in this study. These also show evidence of historical admixture with the southern Arenillas population, suggesting broader past connectivity along the Andean axis. Features such as large geographic extent, ecological breadth, and episodic admixture are generally expected to buffer populations from the accumulation of inbreeding and genetic drift (Chavez et al., 2022; Robinson et al., 2019). However, despite these advantages, we detected the highest burden of long runs of homozygosity in a single individual from Cayambe–Coca (Julio), who notably also exhibited a kinked tail, a phenotype frequently associated with inbreeding in mountain lions (Johnson et al., 2010). If both the elevated ROH burden and the tail malformation reflect recent inbreeding, this pattern may indicate fine-scale population subdivision within the Cayambe–Coca population, with some local subpopulations experiencing recent demographic decline. Importantly, other individuals from Cayambe–Coca did not show comparable ROH signals, suggesting that this effect is not uniform across the region. Together, these observations highlight the need for broader sampling across Cayambe–Coca to test for internal population structure and to determine whether localized declines are occurring in specific areas of this large, protected landscape.

## Conclusions

Our genome-wide analyses demonstrate that Andean puma populations were shaped by regionally contrasting responses to Pleistocene climate dynamics. While populations in the northwestern forests persisted in long-term isolation within stable humid refugia, northern and southern Andean populations experienced episodic connectivity facilitated by transient dispersal corridors during warm interglacial periods. These alternating phases of isolation and contact left distinct genomic signatures that persist in contemporary populations. This historical heterogeneity has direct implications for conservation. Long-term demographic constraints in southern populations have resulted in reduced genetic diversity and elevated deleterious variation, a process that is likely to intensify in the absence of ongoing gene flow. The detection of extensive runs of homozygosity in a single individual from Cayambe–Coca (Julio), despite the absence of a genome-wide pattern of inbreeding, suggests either recent localized inbreeding and population structure in Cayambe-Coca or homozygosity at specific loci associated with developmental traits. By linking climate history with modern genomic vulnerability, this study provides a framework for region-specific conservation strategies. Preserving isolated lineages while restoring or mimicking historical connectivity where appropriate will be essential for maintaining the evolutionary resilience of Andean puma populations.

## Materials and Methods

### DNA Sequencing

To examine the evolutionary history and genomic consequences of demographic change in pumas (Puma concolor), we generated whole-genome sequence data from individuals sampled across ecologically distinct regions of Ecuador, including Andean highlands and coastal forests. Samples were selected to capture geographic and environmental variation within the country. Blood samples from four individuals (Nacho, Mao, Julio, and Ili) were obtained during routine health examinations at the Quito Zoo. Following previous studies in carnivores (Chavez et al., 2026), we took advantage of individuals rescued from human–wildlife conflict and transferred from the wild to zoological facilities; we documented their original localities, which allowed us to incorporate known geographic origins into our analyses. Blood from the remaining two individuals was collected during field sampling in the biological reserve “Mashpi Tayra Wildlife Refuge,” in 2024. All collections were conducted under permits issued by the Ecuadorian Ministerio del Ambiente, Agua y Transición Ecológica (MAE-DNB-CM-2019-0118 and MAATE-DBI-CM-2023-0318).

Genomic DNA was extracted from six puma individuals using the Monarch High Molecular Weight Blood and Tissue DNA Kit (New England Biolabs, USA). DNA quantity and integrity were assessed using a Qubit 2.0 fluorometer (ThermoFisher, USA), a NanoDrop spectrophotometer (ThermoFisher, USA), and agarose gel electrophoresis to confirm high molecular weight DNA prior to library preparation. Sequencing libraries with an average insert size of approximately 150 bp were prepared using the TruSeq Nano PCR-Free kit (Illumina, USA). Whole-genome sequencing was performed on an Illumina NovaSeq X 10B platform, generating 150 bp paired-end reads.

### Read Mapping and Variant Discovery

Raw reads were inspected for quality and processed following an adapted implementation of the Genome Analysis Toolkit (GATK) Best Practices workflow (McKenna et al., 2010). Filtered reads were aligned to the most contiguous chromosome-level Puma concolor reference genome currently available (UCLA assembly; GCA_028749985.4) using BWA-MEM (Li, 2013). The reference genome is approximately 2.5 Gb in length, with an average sequencing coverage of 46×, a contig N50/L50 of 42.3 Mb/17, and a scaffold N50/L50 of 149.8 Mb/7. The assembly is publicly available at https://www.ncbi.nlm.nih.gov/datasets/genome/GCA_028749985.4/.

Duplicate reads were marked, and alignment statistics were evaluated to verify consistent coverage among individuals. Variants were called jointly across all samples using GATK HaplotypeCaller. We retained only high-confidence single nucleotide polymorphisms (SNPs) by applying stringent quality thresholds, including a minimum read depth of five per genotype, Phred-scaled genotype quality scores ≥ 20, and exclusion of sites exceeding the 99th percentile of coverage for each individual to reduce the influence of mapping artifacts. Additional filtering criteria followed GATK recommendations (McKenna et al., 2010) and Chavez et al (2024). CpG sites, insertion–deletion variants, multinucleotide polymorphisms, and loci with more than one alternate allele were excluded prior to downstream analyses. All scripts and command-line parameters are available at https://github.com/dechavezv/2nd.paper.v2.

### Population Structure and Genetic Differentiation

To estimate relatedness among individuals, VCF files were converted to GDS format using SNPRelate (Zheng et al., 2012), enabling compatibility with PLINK-based analyses. Relatedness calculations were conducted on approximately 10,000 high-quality SNPs pruned for linkage disequilibrium (r^2^ ≤ 0.2) and filtered to retain sites with less than 5% missing data. Pairwise kinship coefficients were then estimated using PLINK’s method-of-moments approach for identical-by-descent (IBD) (Purcell et al., 2007), applying a minor allele frequency threshold of 0.05. When kinship coefficients exceeded 0.2, consistent with first-degree relatives, one individual from the pair was randomly removed to avoid bias. This procedure resulted in the removal of one individual from Mashpi.

To assess population structure, we conducted principal component analysis (PCA) using SNPRelate on LD-pruned SNPs (r^2^ ≤ 0.2) filtered at a minor allele frequency of 0.1. Analyses were performed using 500 kb windows across the genome. The first two principal components, which explained the greatest proportion of genetic variance, were visualized using ggplot2 (Wickham, 2016). We further explored ancestry patterns using the Bayesian clustering algorithm fastStructure (Raj et al., 2014). Multiple K values were evaluated, and the optimal number of clusters was determined using marginal likelihood comparisons.

### Evolutionary Relationships and Historic Gene Flow

We used TreeMix (Pickrell & Pritchard, 2012) to infer phylogenetic relationships and test for historical admixture among puma populations. Genome-wide allele frequency data from seven individuals were used to construct a maximum-likelihood population tree. TreeMix models population splits based on allele frequency covariance and then evaluates whether adding migration edges improves the fit between expected and observed covariance patterns.

Analyses were performed on the dataset excluding closely related individuals as identified by SNPRelate (Zheng et al., 2012). We first fitted a strictly bifurcating tree without migration using blocks of 500 SNPs (-k 500) to account for linkage. We then sequentially added migration edges (m = 1 to 10) to evaluate whether models incorporating gene flow better explained the data. Model performance was assessed by comparing likelihood values and examining reductions in residual covariance. A positive migration weight indicated improved model fit when migration edges were included. To determine the best-supported model, we used the -global and -se options to estimate standard errors of residual covariance. Standardized residuals were calculated by dividing deviations between observed and expected covariance by their corresponding standard errors, allowing identification of migration scenarios that most accurately reflected the genetic structure of the populations.

### Genome-wide Diversity

Genome-wide heterozygosity was calculated using non-overlapping 100 kb windows across autosomes for each of the six puma genomes. Heterozygosity was defined as the proportion of heterozygous genotypes relative to all confidently genotyped sites within each window, including homozygous reference and homozygous derived genotypes. Windows with more than 20% missing data were excluded. The script used for these calculations was adapted from Robinson et al. (2019) and is available at https://github.com/dechavezv/2nd.paper.v2/tree/main/4-Demography/Heterozygosity/WindowHet

To quantify genomic signatures of inbreeding, we identified runs of homozygosity (ROH) using PLINK (Purcell et al., 2007). ROH detection employed a sliding-window approach requiring a minimum of 200 SNPs per window, permitting up to three heterozygous calls and 50 missing genotypes. Identified ROH segments were grouped into three size classes: short (< 1 Mb), intermediate (1 – 10 Mb), and long (> 10 Mb). These categories help distinguish signals of ancient demographic contraction from more recent inbreeding events (Ceballos et al., 2018). Total ROH length per individual was expressed as a fraction of the genome.

### Demographic History

To reconstruct long-term effective population size trajectories, we applied the multiple sequentially Markovian coalescent model implemented in MSMC (Schiffels & Durbin, 2014). This method estimates the instantaneous inverse coalescence rate (IICR) through time, which reflects historical changes in effective population size. IICR values were scaled by 2µ to approximate effective population size (Ne), and temporal estimates were converted using a five-year generation time.

### Functional Annotation and Deleterious Variation

To evaluate the accumulation of potentially deleterious mutations in Ecuadorian puma populations, we mapped raw reads to the only fully annotated Puma concolor genome assembly (Saremi et al., 2019). Joint variant calling was performed using GATK HaplotypeCaller v3.8 (Van der Auwera et al., 2013). Functional annotation of variants was conducted using SnpEff (Cingolani et al., 2012) with gene models derived from the genome annotation (~24,000 genes). Variants were classified as synonymous, nonsynonymous (missense), or loss-of-function (e.g., stop-gain or frameshift). When multiple transcripts were available for a gene, the longest transcript was used for annotation. Scripts used for variant annotation and identification of deleterious mutations are available at https://github.com/dechavezv/2nd.paper.v2/tree/main/4-Demography/DeltVariation

### Paleoclimate Reconstruction and Multivariate Environmental Analysis

#### Paleoclimate data and predictor variables

Paleoclimatic variables were obtained from the PaleoClim database, which provides high spatial resolution paleoclimate reconstructions for global terrestrial environments (Brown et al., 2018). Climatic conditions were evaluated across four time periods with a resolution of 2.5 arc-minutes: the Mid-Pliocene Warm Period (mPWP, 3.205 Ma), Marine Isotope Stage 19 (MIS19, ~787 ka), the Last Interglacial (LIG, ~130 ka), and Heinrich Stadial 1 (HS1, 17–14.7 ka).

Environmental predictors included temperature, precipitation, and seasonality variables (BIO1: Annual Mean Temperature, BIO4: Temperature Seasonality, BIO11: Mean Temperature of Coldest Quarter, BIO12: Annual Precipitation, BIO15: Precipitation Seasonality, BIO18: Precipitation of Warmest Quarter), together with elevation (DEM). These variables were selected due to their ecological relevance for structuring climatic gradients in tropical montane ecosystems. All raster layers were spatially harmonized by projecting them into a common coordinate system, aligning raster grids, clipping Ecuador as study extent, and resampling to identical spatial resolution using ArcGIS Pro 3.6.1, as it was for the PCA and climatic distances analyses.

#### Variable standardization

Environmental predictors were standardized using z-transformation:

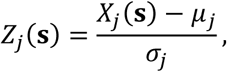

where *X_j_*(*s*) represents the value of predictor *j* at raster cell *s*, and *μ_j_* and *σ_j_* represent the mean and standard deviation of predictor *j*.

Standardization was performed using period-specific statistics for MIS19, LIG, and HS1. In contrast, predictors from the Mid-Pliocene Warm Period were standardized using the statistical moments derived from MIS19. This approach allowed mPWP climatic conditions to be expressed within the same environmental reference framework as MIS19.

#### Principal component analysis (PCA)

Principal component analysis (PCA) was used to summarize climatic variability and reduce multicollinearity among predictors. PCA was conducted using standardized variables following environmental ordination approaches widely applied in ecological niche analyses (Broennimann et al., 2012; Di Cola et al., 2017).

The PCA transformation can be expressed as:

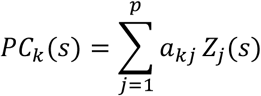

where *a_kj_* represents eigenvector coefficients describing the contribution of predictor *j* to component *k*.

The proportion of explained variance for each component was calculated as:

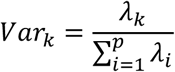

where *λ_k_* represents the eigenvalue associated with component *k*. The first two components (PC1 and PC2) were retained because they captured the dominant climatic gradients within each period.

#### Temporal projection: MIS19-based PCA space for mPWP

To evaluate whether deep-time warm climatic conditions promoted climatic similarity among populations, mPWP climatic predictors were projected onto the MIS19 PCA environmental space. This projection was performed by applying MIS19 eigenvectors to mPWP predictors standardized using MIS19 statistical moments:

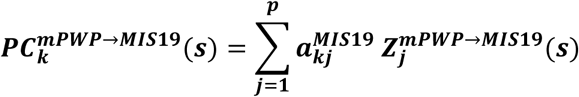

PCA axes depend on the covariance structure of the dataset; therefore, independent PCA analyses across periods produce environmental axes that are not directly comparable. Expressing mPWP climatic conditions within the MIS19 PCA coordinate system enabled direct comparison of climatic similarity across time. This approach follows established methodologies used to evaluate climatic niche stability and environmental analog conditions across temporal scenarios (Broennimann et al., 2012; Di Cola et al., 2017). MIS19 was selected as a baseline because it represents a climatically stable interglacial period that provides a suitable reference for evaluating climatic analog conditions relative to both older and younger climatic intervals.

#### Population location and climatic distances

For each period, we summarized climatic conditions within population polygons (Mashpi, Cayambe-Coca, Arenillas) using zonal statistics on PC1 and PC2 rasters, obtaining mean component scores for each polygon. Climatic dissimilarity between populations *A* and *B* was quantified as Euclidean distance in the two-dimensional PCA space:

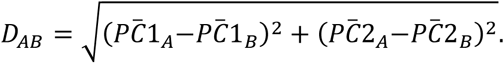

## Data Availability

Raw sequencing reads and accompanying metadata will be archived in GenBank upon publication. Bioinformatic workflows and analysis scripts are available at https://github.com/dechavezv/2nd.paper.v2

## Acknowledgements

We thank Betsabé Trujillo at the Fundación Zoológica del Ecuador for assisting with sample collection. We also thank the staff of the Research and Biology Department of Mashpi Lodge, especially Mateo Roldán; the rangers of the Mashpi–Tayra Wildlife Refuge; and the staff of the Hospital de Fauna Silvestre Tueri (USFQ) for their support on obtaining the samples. We thank Katherine Apunte-Ramos at Pontificia Universidad Católica del Ecuador and Gabriela Pozo at Universidad San Francisco de Quito for assisting with DNA extraction of pumas. We thank Audra Huffmeyer for guidance in data analysis. We are grateful to Brian Jansen, Cindy Hurtado, Álvaro García-Olaechea, and Edward Powers for their contributions. We acknowledge grant FOR-PYI-2.0–0496 from Universidad Tecnológica Indoamérica and funding from Mashpi Lodge, USFQ Fondos COCIBA, and the Ong Family for supporting this research. We collected all samples under permits MAE-DNB-CM-2019-0118 and MAATE-DBI-CM-2023-0318 issued by the Ecuadorian Ministerio del Ambiente, Agua y Transición Ecológica.

## Supplementary Information

### Supplementary Figures

**Figure S1.**
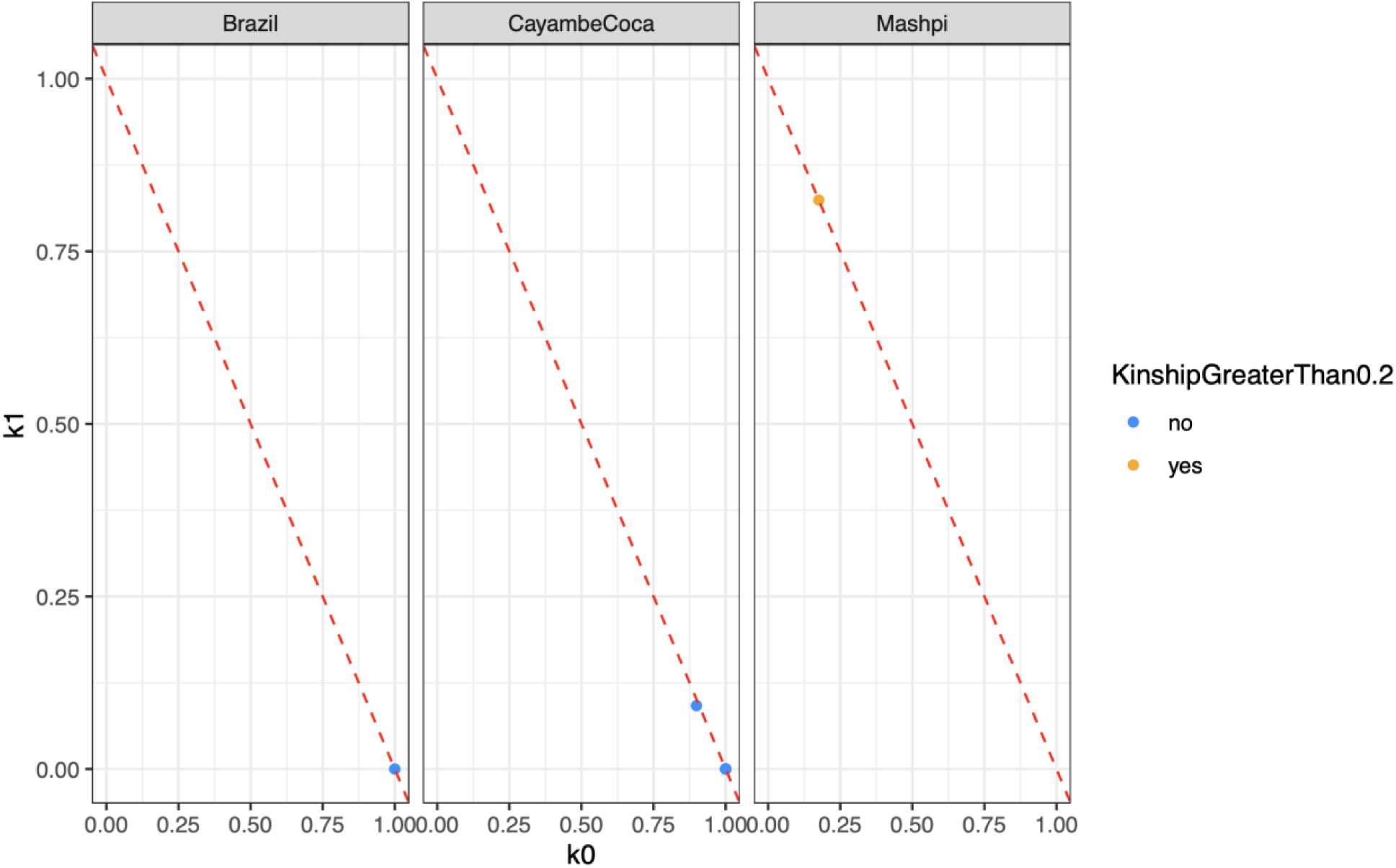
Pairwise kinship coefficients among puma individuals from Brazil, Cayambe–Coca, and Mashpi populations. Scatterplots show estimated kinship values (k0 and k1) for all individual pairs within each population. Points are colored to indicate whether pairwise kinship exceeds 0.2, a threshold commonly used to identify close relatives. The dashed red line represents the expected relationship between k0 and k1 under standard kinship models. This analysis was used to identify and account for close relatives in downstream population genomic analyses.

**Figure S2.**
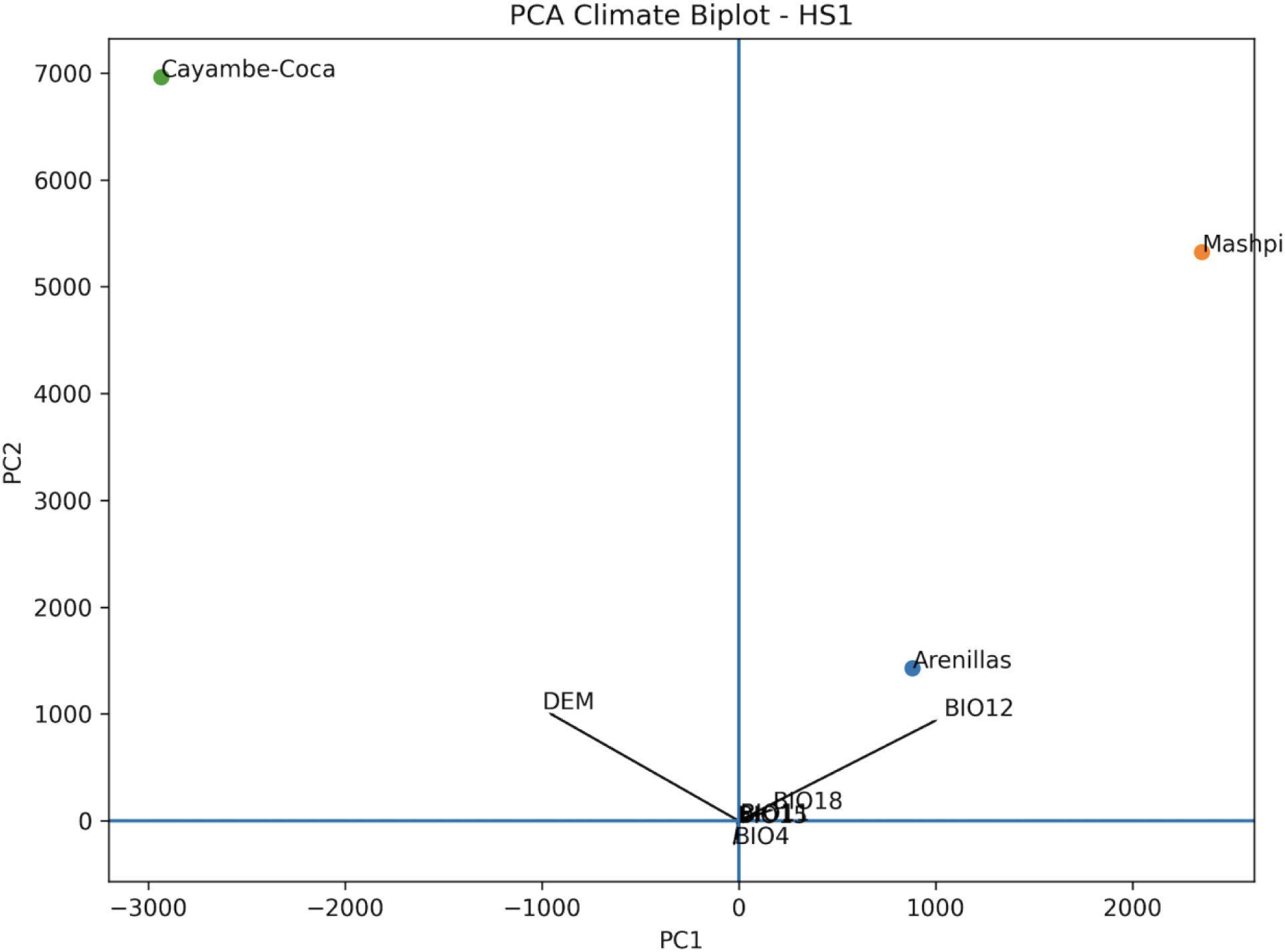
PCA biplots illustrating climatic positioning of Andean Puma populations across the paleoclimatic period Heinrich Stadial 1 (HS1). Arrows represent environmental variable loadings, and points indicate population climatic centroids. HS1 biplots represent period-specific PCA analyses. The first principal component (PC1) primarily represented a humidity–altitude gradient associated with precipitation variables (BIO12 and BIO18) and topographic elevation, whereas the second component (PC2) described a temperature–altitude gradient largely influenced by temperature seasonality and cold-period temperature variables (BIO4 and BIO11).

**Figure S3.**
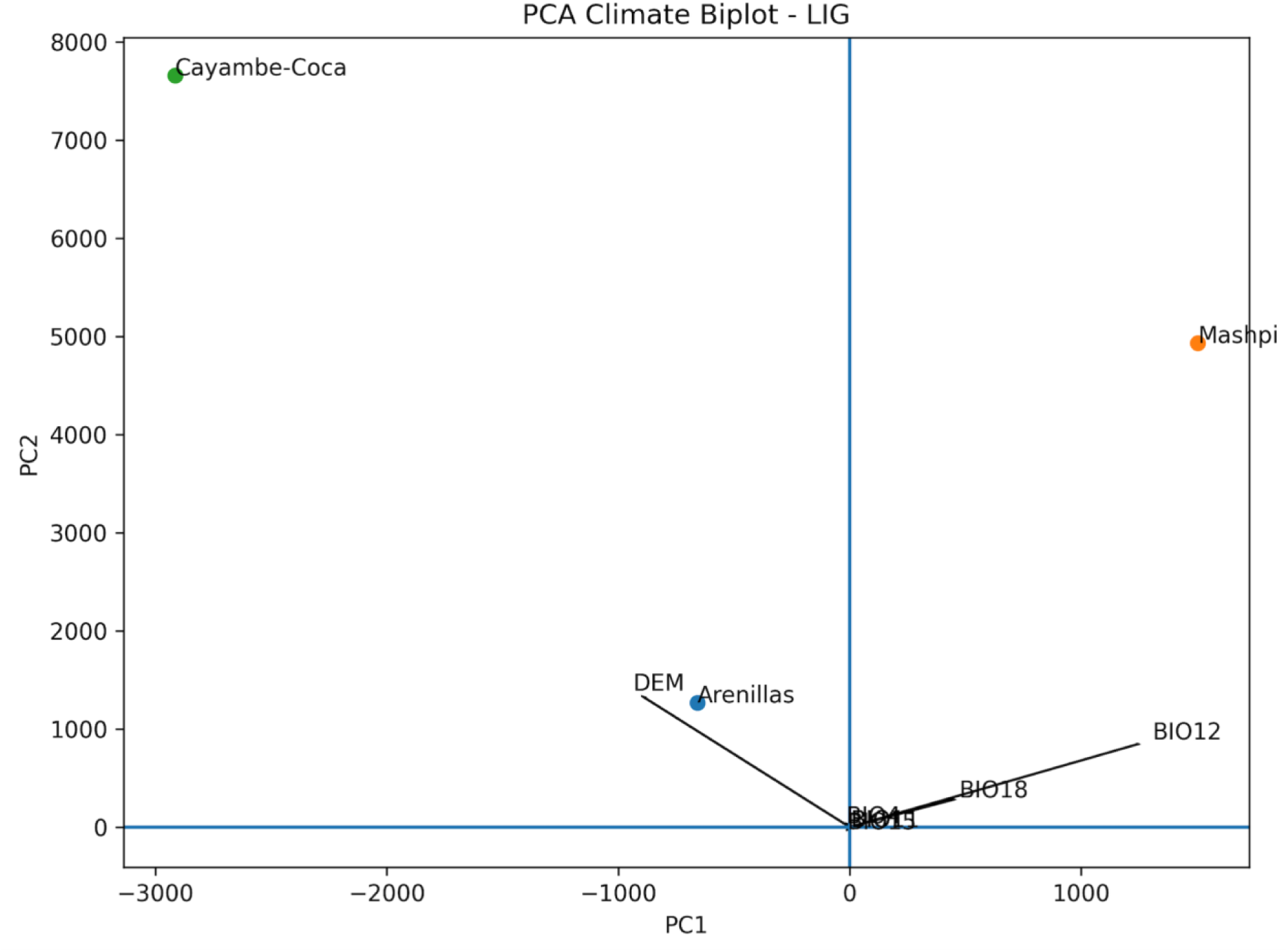
PCA biplots illustrating climatic positioning of Andean Puma populations across Last the Interglacial (LIG). Arrows represent environmental variable loadings, and points indicate population climatic centroids. LIG biplots represent period-specific PCA analyses. The first principal component (PC1) primarily represented a humidity–altitude gradient associated with precipitation variables (BIO12 and BIO18) and topographic elevation, whereas the second component (PC2) described a temperature–altitude gradient largely influenced by temperature seasonality and cold-period temperature variables (BIO4 and BIO11).

**Figure S4.**
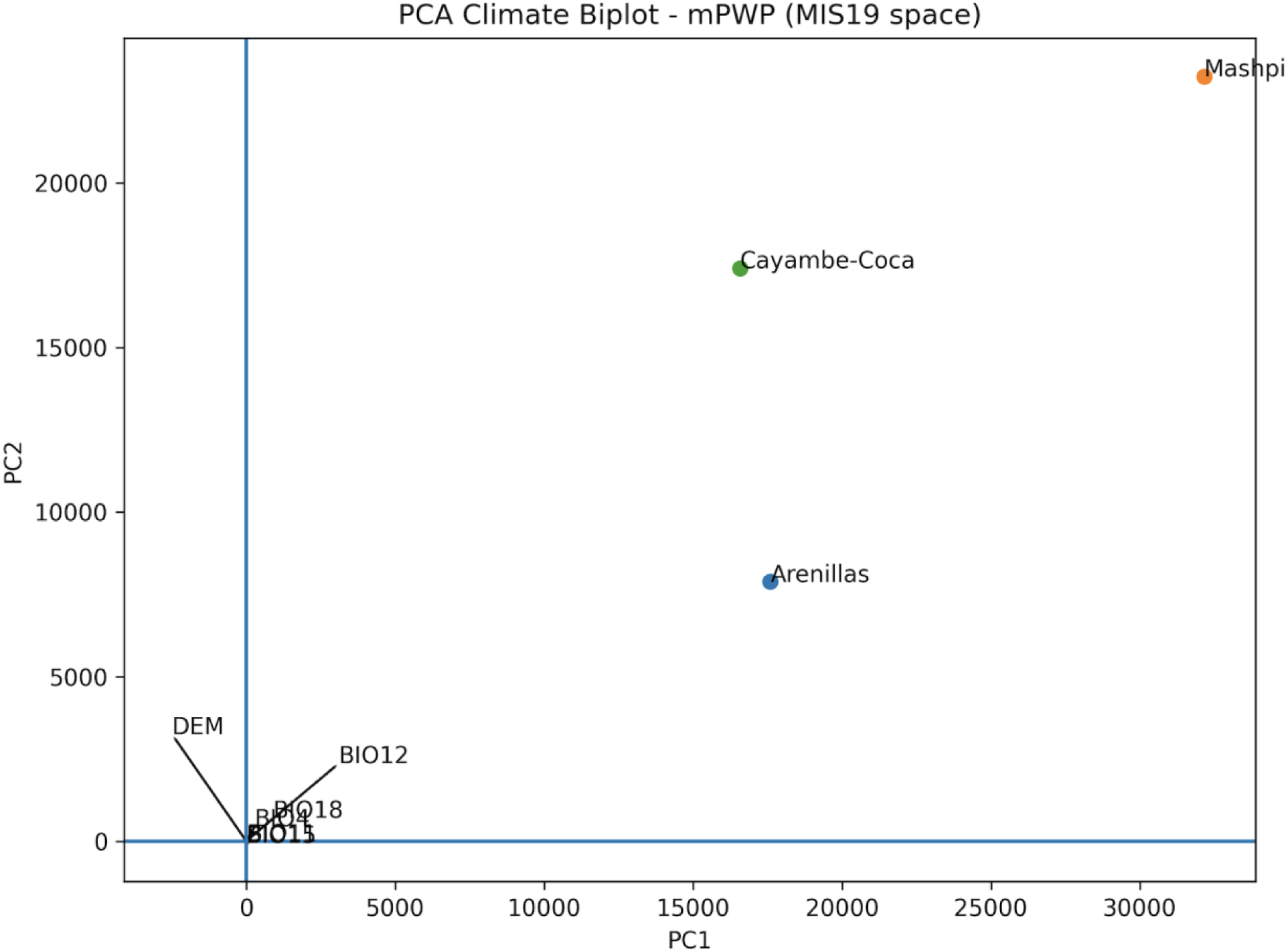
PCA biplots illustrating climatic positioning of Andean Puma populations across Mid-the Pliocene Warm Period. Arrows represent environmental variable loadings, and points indicate population climatic centroids. mPWP climatic conditions are projected into the MIS19 PCA environmental space to allow temporal comparison of climatic similarity with HS1 and LIG. The first principal component (PC1) primarily represented a humidity–altitude gradient associated with precipitation variables (BIO12 and BIO18) and topographic elevation, whereas the second component (PC2) described a temperature–altitude gradient largely influenced by temperature seasonality and cold-period temperature variables (BIO4 and BIO11).

**Supplementary Video link (Kinked tail Julio)** https://docs.google.com/videos/d/15nQvqLNSvccAqosj_vZvOeuwcF92Sj1puySOkxJUlso/edit?u sp=sharing

### Supplementary Tables

**Table S1.** Genome-wide average heterozygosity and total length of short/medium/long runs of homozygosity (ROH) of Ecuadorian pumas.

| Sample ID | Population ID | Hetero(kb*) | Short(MB*) | Medium(MB*) | Long(MB*) |
| --- | --- | --- | --- | --- | --- |
| Ili | Arenillas | 335713 | 162925581 | 146786266 | 21867880 |
| Copal | Mashpi | 426870 | 96878519 | 92472655 | 28581312 |
| Capu | Mashpi | 455981 | 117546237 | 72798886 | 11252821 |
| Nacho | Cayambe-Coca | 475787 | 103596674 | 91127270 | 15898817 |
| Julio | Cayambe-Coca | 484409 | 82824783 | 124598399 | 47030797 |
| Mao | Cayambe-Coca | 490950 | 69314192 | 74955623 | 12226081 |

Note - ROH were categorized as short (<1Mb), medium (1-10Mb), and long (>10Mb).
\* KB stands for kilobases, and MB stands for megabases

**Table S2.** Genotype counts in Andean puma populations.

| SampleId | HomRefCount | HomAltCount | HetCount | MutType |
| --- | --- | --- | --- | --- |
| Mao.CayambeCoca | 1719 | 822 | 873 | LOF |
| Copal.Mashpi | 1833 | 717 | 900 | LOF |
| Capu.Mashpi | 1875 | 765 | 786 | LOF |
| **Ili.Arenillas** | **1839** | **819** | **537** | **LOF** |
| Julio.CayambeCoca | 1818 | 843 | 786 | LOF |
| Nacho.CayambeCoca | 1794 | 777 | 831 | LOF |
| Mao.CayambeCoca | 39850 | 12013 | 18545 | Synon |
| Copal.Mashpi | 40810 | 11441 | 18487 | Synon |
| Capu.Mashpi | 40795 | 11420 | 18594 | Synon |
| Ili.Arenillas | 40756 | 12283 | 14442 | Synon |
| Julio.CayambeCoca | 40013 | 12032 | 18321 | Synon |
| Nacho.CayambeCoca | 40226 | 11945 | 18560 | Synon |
| Mao.CayambeCoca | 27288 | 8457 | 12261 | Missen |
| Copal.Mashpi | 27924 | 8136 | 12500 | Missen |
| Capu.Mashpi | 28005 | 8108 | 12336 | Missen |
| Ili.Arenillas | 27963 | 8279 | 10090 | Missen |
| Julio.CayambeCoca | 27452 | 8662 | 11995 | Missen |
| Nacho.CayambeCoca | 27357 | 8579 | 12376 | Missen |

Note – Homozygote-derived genotypes in the damaging category are higher in populations with smaller effective population sizes, such as the Arenillas population, shown in bold, due to increased drift. In contrast, damaging mutations are less frequent in species with large effective population sizes such as the Cayambe-Coca population. This difference is less evident in the benign category. The counts of benign and damaging alleles in the genome were 17,647 and 2,838, respectively.

### Supplementary Methods

#### ArcGIS Pro workflow

1. **Raster harmonization**

1. Project Raster to a common projected CRS (WGS84).

- Resampling: *Bilinear* (continuous variables).
2. Extract by Mask using the Ecuador boundary polygon.
3. Set Environments consistently for all operations:

- *Snap Raster*: a reference raster.
- *Cell Size*: match the reference raster.
- *Extent*: same as reference raster.
- *Mask*: Ecuador boundary.
4. If temperature layers are stored as °C×10, convert via Raster Calculator.
5. Run Build Raster Statistics on outputs.
2. Standardization to z-scores

- Compute *μ_j_* and *σ_j_* for each predictor raster per period from raster statistics.
- Use Raster Calculator to create Z rasters:

- Period-specific: (X - mean_period) / sd_period for MIS19, LIG, HS1.
- MIS19-based for mPWP: (X_mPWP - mean_MIS19) / sd_MIS19.
- DEM: use a single, consistent DEM_Z (since DEM is time-invariant), aligned to the same grid.
3. PCA rasters

- For MIS19, LIG, HS1:

1. Create a multiband stack with Composite Bands in the fixed band order (BIO1_Z, BIO4_Z, BIO11_Z, BIO12_Z, BIO15_Z, BIO18_Z, DEM_Z).
2. Run Principal Components to generate PCA output rasters and the associated text report containing eigenvalues/eigenvectors.
- For mPWP (projection approach):

1. Do not run Principal Components. Instead, compute *PC*1 and *PC*2 directly with Raster Calculator using the MIS19 eigenvector coefficients (from the MIS19 PCA report):

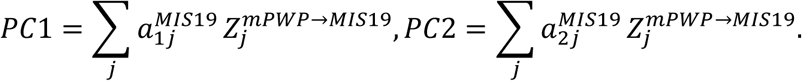
4. Zonal summaries and distances

1. Zonal Statistics as Table for each of PC1 and PC2 using population polygons (zone field = population ID).
2. Join Field to combine PC1 and PC2 summaries.
3. Compute pairwise *D_AB_* distances from polygon means.

## Notes

### Competing Interest Statement

The authors have declared no competing interest.

